# Zinc availability modulates plant growth and immune responses via *AZI1*

**DOI:** 10.1101/166645

**Authors:** Nadia Bouain, Santosh B. Satbhai, Chorpet Saenchai, Guilhem Desbrosses, Pierre Berthomieu, Wolfgang Busch, Hatem Rouached

**Author notes:** Present address: Department of Plant Biology, Carnegie Institution for Science, 260 Panama Street, Stanford, CA 94305, USA. To whom correspondence should be addressed: **Dr. Hatem ROUACHED** UMR Biochimie & Physiologie Moléculaire des Plantes - CNRS-INRA-SUPAGRO-UM, Cedex 2 Montpellier, 34060 France E.mail. **Dr. Wolfgang BUSCH** Salk Institute for Biological Studies 10010 N Torrey Pines Rd · La Jolla, CA 92037 E.mail.

## Abstract

Zinc is an essential micronutrient for all living organisms and is involved in a plethora of processes including growth and development, and immunity. However, it is unknown if there is a common genetic and molecular basis underlying multiple facets of zinc function. Here we used natural variation in *Arabidopsis thaliana* to study the role of zinc in regulating growth. We identify allelic variation of the systemic immunity gene *AZI1* as a key for determining root growth responses to low zinc conditions. We further demonstrate that this gene is important for modulating root growth depending on the zinc and defence status. Finally, we show that the interaction of the immunity signal azelaic acid and zinc level to regulate root growth is conserved in rice. This work demonstrates that there is a common genetic and molecular basis for multiple zinc dependent processes and that nutrient cues can determine the balance of plant growth and immune responses in plants.

## Introduction

Zinc (Zn) is an essential micronutrient for humans, animals, and plants ^1,2,3,4^. It is of particular importance for the function of numerous metalloenzymes that are involved in a plethora of processes such as energy metabolism, nucleic acid and protein synthesis, and protein catabolism^5^. These key biological processes can be adversely altered in situations in which Zn availability is limited. Low Zn manifests itself at physiological and molecular levels, and can cause deleterious effects such as growth retardation and malfunction of immune responses. Recent studies in mammalian systems have shifted the focus on the role of Zn from simply a nutrient to a signalling molecule that fine-tunes intracellular signalling events (for an overview see^6^) and an important player in nutritional immunity^7^. In particular, in the role of Zn for host defence, a complex role is emerging. On one hand, Zn is actively depleted from infection sites restricting the ability of the pathogen to proliferate^8,9^ and on the other hand high Zn levels are generated by the host that contribute to kill the pathogen^10^. However, despite its fundamental importance, it remains unclear whether there is a common molecular basis for these multiple functions involving Zn and whether the signalling and immune related functions of Zn are also relevant for plants.

Plants are the first link in the chain of human Zn nutrition and therefore studying Zn related processes in plants is of particular relevance. Land plants acquire Zn at the root–soil interface and multiple processes are crucial for efficient Zn acquisition. While Zn transport is clearly very important for Zn acquisition and homeostasis^4^, other processes also play pronounced roles for efficient growth under Zn limited conditions. For instance, when responses of two lines of rice that displayed a contrasting tolerance to -Zn were quantified, these lines didn’t differ in Zn-transporter activity but mostly in their maintenance of root growth and the exudation rates of organic acids^11^. A similar correlation of increased root growth and increased tolerance to –Zn conditions had also been described in wheat^12^. Overall, this indicated that Zn levels in the environment are perceived by the plant and lead to distinct changes in root growth that might be important for adaptive responses to low Zn conditions. Nevertheless, so far, neither a role for Zn as signal, nor the genetic and molecular bases of root growth changes upon -Zn conditions in plants has been clearly established yet.

One important biotic stress response process that Zn and its availability has been shown to be involved in animals ^13^ as well as in plants is disease resistance ^14^. One gene that is involved in the response to biotic and abiotic stresses in *Arabidopsis thaliana* is *AZI1* (*AZELAIC ACID INDUCED 1*, At4g12470)^15, 16,17^. It encodes for a lipid transfer protein (LTP)-like protein and belongs to the *EARLY ARABIDOPSIS ALUMINIUM-INDUCED GENE1* (*EARLI1*) gene subfamily^18^. *AZI1* was named after its unique response to the systemically active compound azelaic acid (AzA, a nine-carbon dicarboxylic acid)^19^. In the Arabidopsis genome, *AZI1* clusters in a tandem array on chromosome 4 with three other *EARLI1*-type genes, namely At4g12480 (*EARLI1*), AT4G12490 (*ELHYPRP2 (EARLI1-LIKE HYBRID PROLINE-RICH PROTEIN 2*) and AT4G12500. Among these genes, the role of *AZI1* for long-distance signals related to systemic acquired resistance (SAR) is the best documented so far. In addition to *AZI1*, SAR involves an important hormone, namely salicylic acid (SA). SA accumulates upon pathogen attack^20^, and leads to the induction of *PATHOGENESIS-RELATED GENE 1* (*PR1*, SA marker gene) ^21^. In *Arabidopsis thaliana*, mutation of *AZI1* causes a specific loss of systemic immunity triggered by pathogens^19^. Beyond its role in biotic stress responses, the involvement of *AZI1* in response to abiotic stress, such as the regulation of seedling growth under salt stress, was demonstrated^17^. But, whether *AZI1* is involved in the response to nutrient levels that potentially affect the plants capability to defend itself or the capability for pathogens for infection (e.g. low Zn), remains unexplored.

Here, we approach study the genetic basis of low exogenous Zn levels on root growth by exploring natural genetic variation. We find that there is heritable natural variation of root growth responses to low Zn and that natural allelic variation of the immune gene *AZELAIC ACID INDUCED* (*AZI1*) determines a significant proportion of this response. We further reveal an intriguing evolutionary conserved interaction between exogenous Zn levels and AzA dependent defence pathways to regulate root growth.

## Results

### The AZI1 gene is involved in the root growth response to low Zn conditions

To identify genetic components that regulate plant growth upon low Zn (-Zn) conditions, we determined the length of the primary root of 230 genetically diverse natural accessions of *Arabidopsis thaliana* (Figure S1A) from the RegMap population^22^ grown on +Zn and -Zn medium over 7 days (Table S1-2). Importantly, while still being correlated, the root growth of the vast majority of accessions clearly differed in +Zn and –Zn conditions (Figure S1B, C). To assess whether these root growth responses were specifically due to the -Zn treatments, we determined mRNA levels of four Zn-deficiency responsive marker genes *ZIP3*, *ZIP5*, *ZIP12* ^23^ and *PHO1;H3*^24^ in the Col-0 accession under our screening conditions. All of these four genes were significantly up-regulated in –Zn conditions (Figure S2), demonstrating that the plants sensed and responded to the -Zn conditions.

In the panel of screened accessions, we observed broad phenotypic variation for root length (Figure S3) that was highly heritable (broad sense heritability (H^2^) ranging^25^ from 0.36 to 0.44) (Table S3). We then conducted Genome Wide Association Studies (GWASs) using the AMM method that corrects for population structure confounding^26^, to identify loci that were associated with root length under -Zn (Figure 1A, Figure 1B, Figure S4 and Figure S5). We then corrected the association *P*-values for all SNPs for multiple testing using the Benjamini-Hochberg-Yekutieli method^27^. Due to the limited power of our association study and the potentially high false negative rate due to population structure correction, and because we aimed to test any major emerging candidate experimentally, we selected a relatively non-conservative 10% false discovery rate (FDR) as our threshold for significant association (we thus expect that 10% of the significant SNPs are false positives). Using this criterion, we identified two chromosomal regions associated with root length in –Zn conditions. On chromosome 2, the significant peak (*P*-value = 3.27*10^-7^; FDR ≈ 7%) was located in a region with a cluster of similar genes encoding a Cysteine/Histidine-rich C1 domain family (At2g21810). It was detected on the last day of the time course (day 7). These proteins require Zn ions for their function^28^ (pfam, PF00130). On chromosome 4, the significant peak (*P*-value = 4.40*10^-7^; FDR ≈ 6%) was located in in a region that contains the lipid transfer protein (LTP)-like *AZI1* (At4g12470) gene and the 7 additional genes encoding for lipid transfer proteins as a cluster (Figure 1B and Figure 1C). This peak was already detected early in the duration of our time course (day 2) suggesting that this locus was relevant for Zn signaling rather than being a consequence of low internal Zn levels. As *AZI1* itself was known for being involved in signalling: it mediates azelaic-acid-induced systemic immunity^19^; we hypothesized that *AZI1* was involved in mediating crosstalk between nutrient and immunity signals. However, as the GWAS peak spanned multiple genes, we first tested whether the best candidate in this region was indeed *AZI1*. For this, we assessed all 8 genes in the genomic region surrounding the association peak. Of these 8 genes, only *AZI1* showed significant transcript level alteration in response to -Zn in Col-0 (Figure 1D), providing a strong hint that it was involved in –Zn dependent root growth regulation. To test this further, we determined the root lengths of Arabidopsis Col-0 (WT), *azi1* (T-DNA) mutant line and an *AZI1* overexpressing (OE *AZI1*) line (35S::*AZI1*) grown in +Zn or -Zn conditions over 7 days (Figure S6). In presence of Zn, no significant differences in root length could be observed between the *azi1* mutant and wild-type plants (day2, Figure 1E; 7 days Figure S6). Grown under -Zn, the root length of *azi1* was significantly shorter than Col-0 and 35S::*AZI1* plants (day 2, Figure 1E, 7 days, Figure S6). Interestingly, starting from day 5 onwards, compared to the Col-0 wildtype, roots of *azi1* were significantly shorter and roots of 35S::*AZI1* plants were significantly longer respectively (Figure S6; day 5, Figure S7 A-B), which suggests that the expression level of *AZI1* is involved in controlling this trait (root growth). Day 5 was therefore chosen as time point for further analysis. To assess whether this is a function of *AZI1* that is common to other micronutrients, we grew the same set of lines under low iron (-Fe) conditions (Figure S7-C). There, no significant root length difference could be detected for the 3 tested lines, which indicates that growth responses to comparable nutrient limitations are not dependent on *AZI1* and supports the notion of a rather specific *AZI1* dependent response to Zn. Taken together, these data show that *AZI1*, previously described as a key components of plant systemic immunity involved in priming defence^19,29^, modulates root growth in a Zn level dependent manner.

**Figure 1.**
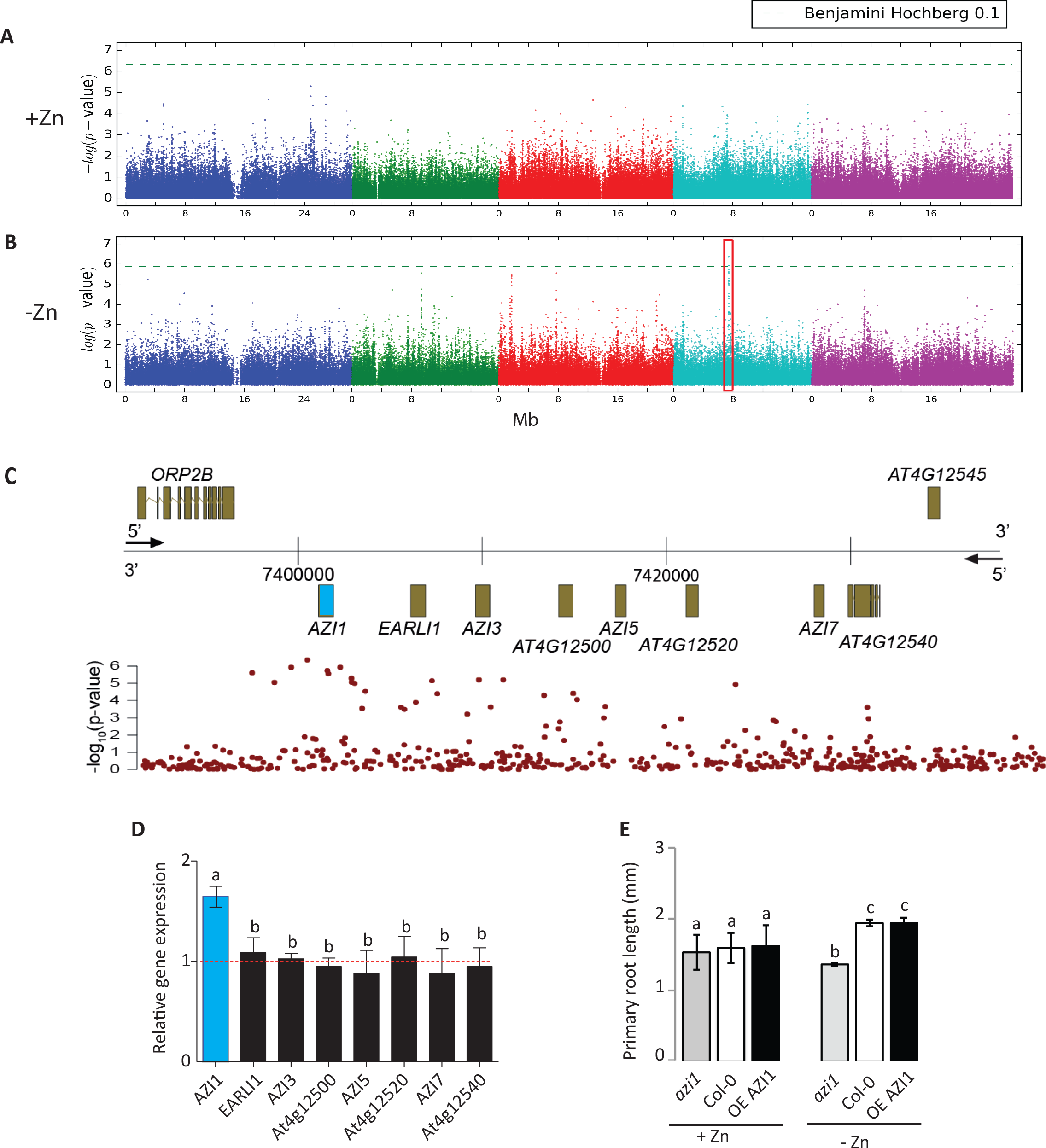
AZI1 controls root growth under zinc limiting conditions. GWAS for mean root length based on a set of 230 *A. thaliana* accessions grown under (A) zinc sufficiency (+Zn) or (B) low zinc (-Zn) (day 2). The chromosomes are represented in different colors. The horizontal dash-dot line corresponds to a FDR of 10% after Benjamini–Hochberg–Yekutieli correction. The red box indicates the significant association. (C) Genomic region around *AZI1* locus (highlighted in blue). X-axis: genomic position. Y-axis: upper panel: Gene models. Lower panel: LOD score from GWAS depicted in (B). (D) Expression changes (fold change) of *AZI1*, *EARLI1*, AZI3, At4g12500, AZI5, At4g12520, *AZI7* and At4g12540 in Col-0 grown upon growth in –Zn conditions compared to Col-0 plants grown under +Zn conditions. Every data point was obtained from the analysis of roots collected from a pool of ten plants. Error bars correspond to s.d.; three biological repeats. The *Ubiquitin* gene was used as an internal reference. (E) Average primary root length (Day 2) of wild-type plants (Col-0 genotype), *azi1* mutant and overexpressor line 35S::*AZI1* (OE *AZI1*) plants grown under +Zn or –Zn respectively. Experiments were independently repeated three times, and data are represented as mean ± s.d. n = 10. Letters a and b indicate significantly different values at p <0.05 determined by one-way ANOVA and Tukey HSD.

### Natural allelic variation of AZI1 determines root growth under zinc limiting conditions

While we had shown that *AZI1* was involved in modulating root growth in a Zn level dependent manner, this was no proof that the allelic variation of *AZI1* is causal for the observed root growth differences under –Zn. We therefore set out to test this. Sequence analysis of the *AZI1* genomic region (promoter and coding region) showed multiple polymorphisms in the regulatory region as well as synonymous changes in the coding region (Table S4) that were consistent different between contrasting groups of accessions with either long or short roots on –Zn (Figure S8A and Figure S8B, table S1 and table S2). Consistent with causal regulatory polymorphisms, *AZI1* expression was significantly higher upon –Zn in accessions with longer roots (Figure S8C and Figure S8D). For further analysis, we focussed on two contrasting accessions, Col-0 and Sq-1, which were among the most contrasting accessions regarding their root length on –Zn (Table S1 and Table S2) and each displayed the variant of the marker SNP that was associated with long and short roots on –Zn respectively. To then experimentally test whether the difference in *AZI1* expression level was due to the natural allelic variation and whether this was also causal for the longer roots, we transformed the *azi1* mutant (Col-0 background) with constructs containing 1.6kbp of the promoter and the coding region from either Col-0 (long roots in -Zn) or Sq-1 (short roots in -Zn), and an empty vector (control). In five independent homozygous single insertion lines (Table S5) complemented with the Col-0 pAZI1:*AZI1* the expression level of *AZI1* under –Zn was significantly higher (P<0.01) than that in plants transformed with the Sq-1 pAZI1:*AZI1* construct (Figure 2A). Consequently, we tested these T3 lines for root length differences under –Zn and +Zn. Consistent with the hypothesis that our *AZI1* variants determine root growth specifically under –Zn, no difference in term of root length was observed between the T3 lines grown on +Zn (Figure S9), while under *-Zn*, we observed significantly longer roots (P< 0.05) in the Col-0 pAZI1:*AZI1* plants compared to Sq-1 pAZI1:*AZI1* plants or *azi1* plants transformed with the empty vector (Figure 2B). Taken together, these data demonstrate that allelic variation at the *AZI1* locus causes variation of *AZI1* expression levels and that this leads to variation of PRG under -Zn. As there were multiple polymorphisms in the regulatory region of the *AZI1* gene (Col-0 and Sq-1 accessions, Figure S10), and only synonymous changes in the coding sequence in these constructs, the data also suggest that this regulation is mainly dependent on regulatory changes. We note, that we can’t completely exclude the additional involvement of other genes in the associated region in contributing to this response.

**Figure 2.**
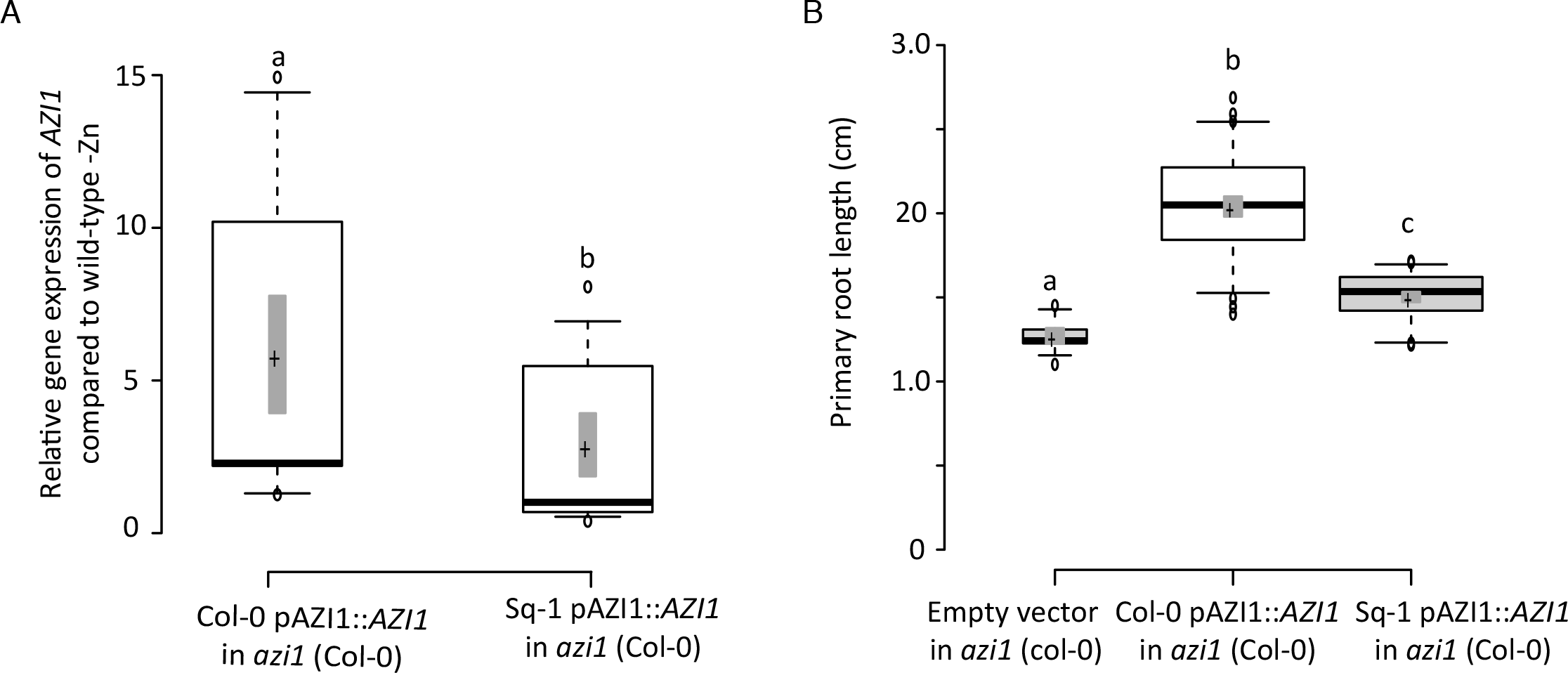
Natural allelic variation of the AZI1 locus underlies phenotypic variation of root length in zinc limiting conditions. (A) Transcript level of *AZI1* in *azi1* lines complemented with pAZI1:*AZI* from either Col-0 or Sq-1 on –Zn conditions and shown as relative to +Zn condition (5 days). Five independent T3 lines were considered for this analysis. Relative expression was quantified in three biological replicates using RT-qPCR. The Ubiquitin gene was used as an internal reference. (B) Primary root length of *azi1* lines complemented with pAZI1*:AZI1* from either Col-0 (n=50), Sq-1 (n=50) or the empty vector (n=10) on –Zn conditions (5 days). For each genotype, three repeats each containing five independent T3 lines. Box plots show analysis of the relative gene expression of *AZI1* (A) and primary root length (B) in of *azi1* lines complemented with pAZI1:*AZI* from either Col-0, Sq-1 or the empty vector on –Zn conditions. Center lines show the medians; box limits indicate the 25th and 75th percentiles as determined by R software; whiskers extend 1.5 times the interquartile range from the 25th and 75th percentiles. Letters a, b and c indicate significantly different values at p <0.05 determined by one-way ANOVA and Tukey HSD.

### Azelaic Acid Inhibits Arabidopsis Root Growth in a Zn-dependant manner

While *AZI1* had not been implicated in any known process involving Zn, it is known to mediate signal mobilization for systemic defence priming that can be triggered by AzA^19,20^. We therefore hypothesized that *AZI1* would modulate growth and immunity programs depending on Zn and AzA status. To test this hypothesis, we first established the effects of the exogenous application of AzA on root growth. AzA affected root growth in a dose-dependent manner starting with a relatively mild reduction of growth at 100 µM to complete inhibition of root growth at 200 µM AzA (Figure 3A). We then determined whether this response is dependent on *AZI1*, and assessed root growth in Col-0 and *azi1* mutant at 100 µM AzA and in presence or absence of Zn after 5 d of treatment. While, AzA severely inhibited root growth in Col-0 plants in presence of AzA and Zn (+AzA+Zn), the *azi1* mutant line was significantly more resistant to the inhibitory effect of AzA (Figure 3B). This demonstrated that AzA modulates root growth in an *AZI1* dependent manner.

**Figure 3.**
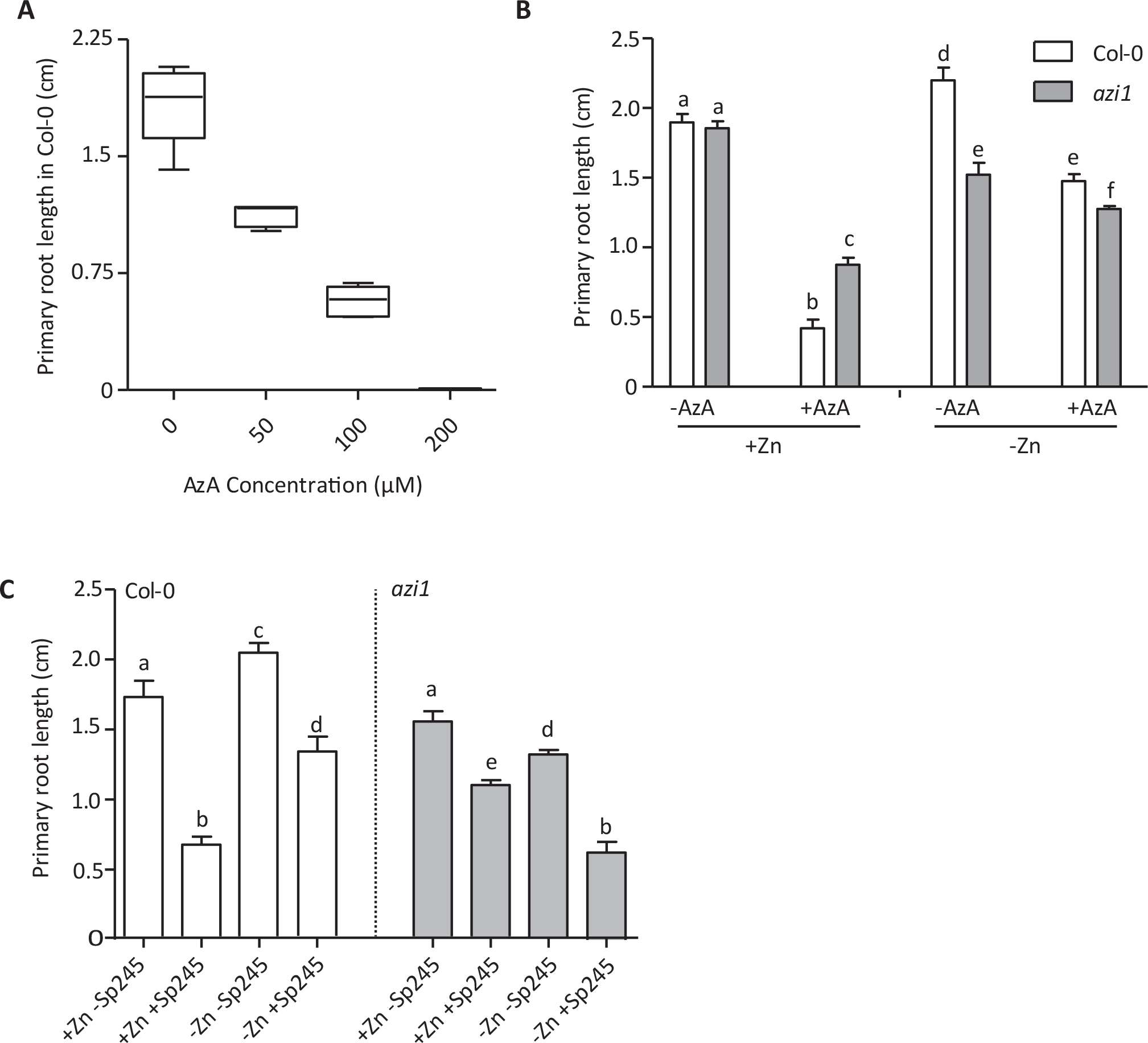
Azelaic acid, zinc level and bacterial presence interact to regulate root length and in an AZI1 dependent fashion. (A) Root lengths of wild-type seedlings treated with different AzA concentrations on +Zn or –Zn medium for five days. Box and whisker plots were generated using Prism (Graphpad), with the box represents the 25th to 75th percentiles and the whiskers reaching to the lowest and highest values. The line in the box shows the median. (B) Root lengths of 5-day-old Col-0 and *azi1* seedlings treated with 100µM AzA on +Zn or –Zn medium. (C) Root lengths of Col-0 and *azi1* seedlings grown with *Azospirillum brasilense* (Sp245) on +Zn or –Zn medium for five days. Data are represented as mean ± s.d. (n = 10). Letters indicate significantly different values at p <0.05 determined by one-way ANOVA and Tukey HSD.

To test whether Zn modulates this response, we conducted the same assays under –Zn conditions. Strikingly, low levels of Zn alleviated the growth inhibitory effect of AzA on Col-0 to a large extent, and led to a further increased growth of the *azi1* mutant line (Figure 3B). Taken together, these data show that AzA induced reduction of growth is modulated by Zn levels, and that *AZI1* is a key component for this modulation.

### Zn status strongly impacts immunity and modulates the response to AzA

Our data had not only shown that Zn levels and AzA modulate root growth, but also that the root growth responses to these treatments strongly interact (Figure 3B). To test whether this interaction is due to the modulation of molecular responses to AzA by Zn levels, we measured the expression levels of 18 defence-related genes that had been shown a mild but significant expression change upon AzA treatment (P < 0.05) in leaves of wild-type plants (Col-0) ^19^, as well as 2 additional genes regulating salicylic acid biosynthesis (*WRKY28* and *WRKY46)* ^30,31,32^, *AZI1*, and a marker gene frequently used as a reliable molecular **marker** for **SA**-dependent SAR (*PR1*)^21^. Our q-RTPCR based gene expression analysis showed that almost all (16) of these defence-related genes were upregulated in response to the application of the application of AzA (Figure S11). Notably, the group of most strongly induced genes contained genes involved in salicylic acid (SA) biosynthesis, such as *ISOCHORISMATE SYNTHASE 1*, *WRKY28*, *WRKY46*, as well as the SA response marker *PR1*. AzA treatment of plants grown on –Zn medium (-Zn+AzA) resulted the upregulation of only 9 of the 16 genes that were upregulated in +Zn/+AzA (Figure S11). Notably, the SA response marker gene *PR1* was not among these. Overall this suggests that interaction of –Zn and AzA is not due to a lack of induction of SA biosynthesis genes, but rather acts more downstream during SA signalling. Furthermore, consistent with an effect of Zn levels on the expression of these genes, plants grown on low Zn showed a down-regulation of 6 of the 16 defence-related genes induced by AzA alone (Figure S11). Overall, these results demonstrate that Zn status impacts the expression of defence-related genes and modulates the response to AzA in plants.

### A. brasilense Infection Inhibits Root Growth in Arabidopsis in a Zn-dependant manner

The link of Zn status and *AZI1* to both, growth and defence prompted us to test whether interaction of *AZI1* and Zn status impacts growth responses to biotic stimuli. For this, we chose to measure primary root responses upon infection with the bacterium *Azospirillum brasilense* (Sp245) as it had been described that root growth as well as *AZI1* expression is changed in *A. thaliana* upon such infection^33^. Consistently with^33^ root growth of 5 d old WT plants was reduced when grown on complete medium (+Zn) and inoculated with *A. brasilense* (Figure 4A). When grown on –Zn, this *A. brasilense* induced root length reduction was slightly less pronounced (Figure 3C). The bacterially induced growth inhibition under +Zn was significantly less pronounced in the *azi1* mutant, showing that *AZI1* is involved in regulating this growth response. Under –Zn, bacterial incubation resulted in further decrease of root growth of *azi1* mutant compared to –Zn (Figure 3C). Overall, these data clearly show that there is a complex interaction of Zn levels and the *AZI1* gene that determines the balance of growth and defence.

**Figure 4.**
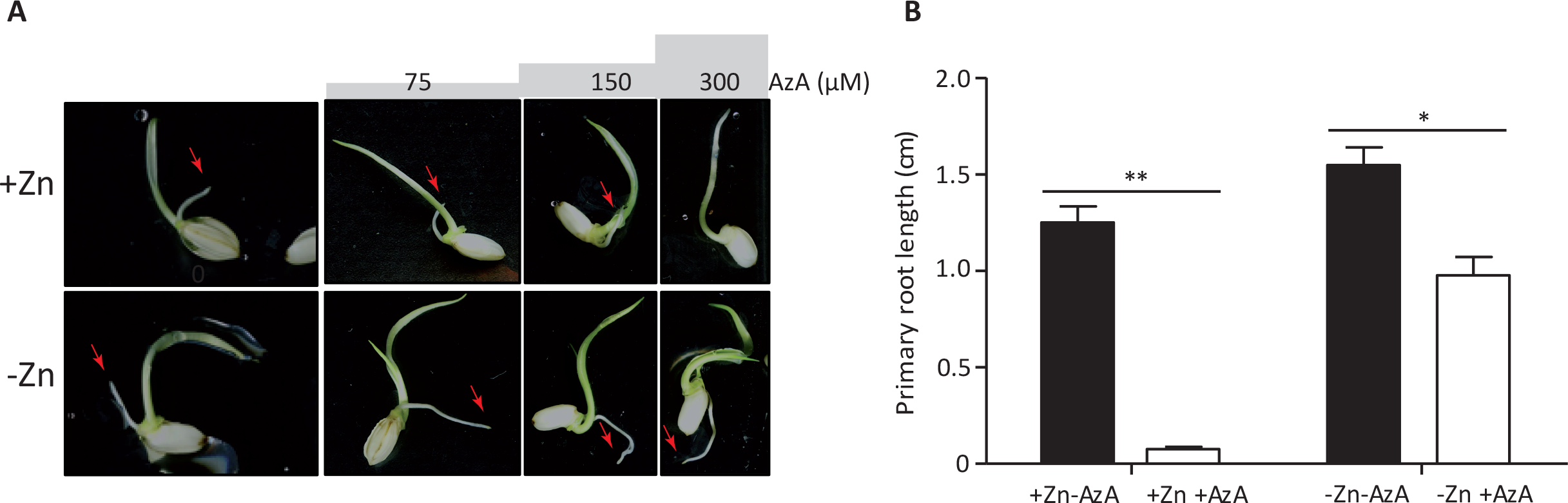
Azelaic acid and zinc interact to regulate root growth in Oryza sativa. A) Images of representative Rice (*Oryza sativa*, Nipponbare) seedlings grown on Yoshida medium supplemented with 300 µM of AzA presence or absence of Zn for five days. Red arrow indicates root. (B) Quantified primary root length of Rice seedlings (n=10). The data are given as means ± s.d. Asterisk indicates statistical significance, P < 0.05. Double asterisk indicates statistical significance, P < 0.01.

### Zn and AzA interaction is conserved in Oryza sativa root

The gene family to which *AZI1* belongs, is strongly conserved throughout the *Viridiplantae* (green plants) ^34^. This led us to hypothesize that the AzA mediated root growth regulation and its modulation by Zn status is a conserved growth-immunity regulating pathway. We therefore investigated the effect of AzA on root growth in presence and absence of Zn in the monocot species rice (*Oryza sativa*) (Figure 4A and Figure 4B). Root growth increases in rice grown in low Zn compared to +Zn condition (Figure 4A and Figure 4B). Interestingly, while seedlings grown in medium that contained AzA (300 µM) and Zn didn’t develop any roots, this growth inhibition was not observed when germinating the grains on –Zn medium containing AzA (Figure 4, Figure S12). This demonstrated that AzA mediated growth inhibition, as well as its regulation by Zn levels is an evolutionary conserved mechanism.

## Discussion

Plants must sense changes in external and internal mineral nutrient concentrations and adjust growth to match resource availability ^35^. Responses to nutrient limitations manifest very early in plants life cycle, and root related processes are a major target of responses to nutrient constraints in plants. Consequently, primary root growth responds early on and drastically to nutrient limitations^36^ and genome wide association mapping approaches are now being used to understand the genetic and molecular factors that govern these early growth responses^37^. However, apart from these abiotic variables (e.g. nutrients and water levels), the root continuously is exposed to changing biotic factors. Our study to identify genes and their variants that determine early root growth responses to low Zn levels revealed a genetic and molecular between root growth response to abiotic (Zn) and biotic (defence related signalling) factors. In particular, our study revealed a significant association between the *AZI1* locus and primary root length in Arabidopsis accessions grown in –Zn condition, as well as an intriguing *AZI1* dependent interaction between Zn levels and the AzA pathway. One interesting question is whether the *AZI1* function that relates to abiotic factors are specific to Zn or not. While there has been one report in which it was shown that ectopic expression of *AZI1* improved Col-0 seed germination under high salinity stress condition and *azi1* mutants were overly sensitive^17^, many leads point in the direction of a largely Zn specific function: Significant associations close to the the *AZI1* locus were neither found in the GWAS for accessions that were grown at the same time on +Zn (Figure S2, Figure S3, Figure S4 and Figure S5), nor in other published root GWAS datasets for growth on MS medium^38^, under different nitrogen conditions^39^, under NaCl stress^40^, and under Fe deficiency^37^. Moreover, while loss of function of *AZI1* caused decreased root growth of Arabidopsis plants grown under –Zn conditions, and *AZI1* overexpression caused the roots to be longer than WT (Figure 1), this was not observed when *azi1* or *35S:AZI1* were grown under -Fe conditions (Figure S6). Nevertheless, the level of exclusiveness or the extent of this specificity of *AZI1* function for Zn can however only be elucidated by further studies.

The responses that we have observed and studied occur early-on during plant development, when Zn levels in the seedlings will not be significantly depleted. Moreover, it is likely that traces of Zn will have remained in the washed agar that we used for these assays, generating an environment very low in Zn, rather than a fully Zn depleted growth environment. Overall, our findings thus exposed mechanisms that will relate more to sensing than to a Zn starvation/depletion response, consistent with our observation that these low Zn levels promote early root growth rather than to inhibit it. While we have identified a signalling mechanism that contributes to this –Zn dependent increase in root growth, the Zn sensor still remains elusive.

Our study proposes a possible mechanism for the regulation of root growth depending on the environmental Zn level, in which AZI1 plays an important role, and probably in an interaction with SA. AZI1-and SA-related signals are known to interact^20^, and possibly involved in a self-amplifying feedback loop (for review^41,42^). Our gene expression analysis revealed that Zn status impacts the expression of *AZI1*, as well as other immune-and SA-related genes in Arabidopsis (Figure S11). Variation in SA concentration in leaves and roots of Arabidopsis plants upon nutrient deficiency stresses has been reported^43^, and appear to be nutrient specific^43^. For example, SA levels significantly increased in response to potassium deficiency while low Fe caused a significant decrease of SA in roots^43^. The involvement of phytohormones in controlling root growth under different nutritional status has been documented^44^, and hormones accumulation “threshold” appeared to be critical for hormones action^45^. For example, in various plants species, treatment with low concentrations of SA led to root growth promotion while treatment with high concentration of SA caused an inhibition of root growth^41,42^. Plants exposed to bacterial infection accumulate SA^20^, AzA, and induce the expression of *AZI1*^33^ presumably to promote defence response, often at the cost of growth. In our gene expression analysis, a marker frequently used for SA accumulation, namely *PR1*^21^, was induced by +Zn+AzA conditions, suggesting an accumulation of SA under these conditions. Consistently, root growth of wild-type plants (Col-0) treated with (+Zn+AzA) was severely reduced compared to control condition (+Zn-AzA). Similarly, plants (Col-0) treated with Sp245 showed shorter root than the ones grown on control condition. The negative effect of either AzA or Sp245 treatment on root growth was alleviated when these stresses were combined with low Zn treatment, (-Zn+AzA or -Zn+Sp245). This was accompanied by an absence of *PR1* induction, indicating that the otherwise root growth inhibitory SA response is affected by low Zn levels at the molecular level. Our work exposed an interaction of Zn levels and immunity and a common genetic and molecular basis for this. Interestingly it seems that Arabidopsis can prioritize root growth over defence responses, if Zn levels are low during early development. The data from rice point in the same direction and provide a hint that this interaction is evolutionary conserved. It will be interesting to elucidate how this relates to the molecular role of Zn in nutritional immunity in plants, perhaps somehow similar to the role of Zn levels in infection sites in animal systems^8,9,10^.

*AZI1* belongs to a large family of pathogenesis-related proteins; LTPs. While some LTPs were suggested to play a role in defence reaction in a root specific manner^46^, the role for AZI1 in defence responses was mainly investigated in aboveground tissues, and similar data for role of *AZI1* in roots are still absent. In addition to *AZI1*, roles of AzA had been analysed on the aboveground tissues, our data reveal an important role of this pathway in the root, which extends the current models for underground defence priming. Importantly, the conserved specific interaction of Zn and AzA that can be observed not only in the dicot species Arabidopsis but also in the monocot species rice, is not only interesting from the view of basic biology, but also harbours very interesting perspectives for an innovative biotechnological application. This is because AzA is thought to directly mediate crop plant responses to pathogens and herbivores or to mimic compounds that do^47^ and is listed among the natural compounds that induce resistance by a priming mechanism^48^. To activate these plant responses, AzA like others organic and inorganic chemicals can be applied as a foliar spray, seed treatment, or soil drench^47^. However, our work revealed that the soil-based application of AzA might significantly impact root traits depending on Zn bioavailability (AzA severely inhibited the root growth in +Zn), which could have an enormous and direct impact on plant growth in the field.

## Acknowledgments

The authors are thankful to Prof. Peter Urwin (University of Leeds, UK) for providing seeds of *azi1* mutant and Dr. Youssef Belkhadir, Dr. Hironaka Tsukagoshi and Dr. Zaigham Shahzad for helpful discussions and suggestions. We are also grateful to Bonnie Wohlrab, Christian Göschl and Mushtak Kisko for technical assistance. This work was funded by the “Institut National de la Recherche Agronomique – Montpellier - France” INRA (H.R.), and supported by funds from the Austrian Academy of Science through the Gregor Mendel Institute (GMI) and an Austrian Science Fund (FWF) stand-alone project (P27163-B22) (W.B.).

## Author information

### Contributions

H.R and W.B. designed the research. H.R supervised this project. N.B, S.B.S and C.S. performed most experiments. G.D. helped conduct bacterial infection experiments. P.B. helped conducting the Zn root phenotyping data analysis. H.R and W.B. analyzed the data and wrote the manuscript.

### Corresponding authors

Correspondence to: Hatem Rouached or Wolfgang Busch

## Material and Methods

### Plant materials and growth conditions

We used 230 different genotype of *A. thaliana* from different geographic origins (Table S1, S2, Figure S1A) and for each genotype grew 12 seedlings. The previously described ^49^ *azi1* insertion mutant (SALK_085727C) available in the Nottingham Arabidopsis Stock Centre was provided by Peter Urwin (University of Leeds, UK). Lines overexpressing *AZI1* under 35S promoter, expressing Col-0 or Sq-1 *AZI1* locus under their native promoters were generated in this *azi1* mutant background. Plant phenotyping for GWAS was as described previously^38^. Briefly, for each growth condition all lines were grown side by side. For both Zn conditions, GWAS assays were performed in the same growth chambers under the same 22°C long-day conditions (16 h light, 8 h dark). Seeds were placed for 1 □h in opened 1.5 □ ml Eppendorf tubes in a sealed box containing chlorine gas generated from 130 □ ml of 10% sodium hypochlorite and 3.5□ ml of 37% hydrochloric acid. For stratification, seeds were imbibed in water and stratified in the dark at 4 □°C for 3 days to promote uniform germination. On each plate, eight different accessions with three seeds per accession were then germinated and grown in a vertical position on agar-solidified medium contained 0.5 mM KNO3, 1 mM MgSO4, 1 mM KH2PO4, 0.25 mM Ca(NO3)2, l00 μM NaFeEDTA, 30 μ M H3BO3, l0 μM MnCl2, l μM CuCl2, 15 μM ZnSO4, 0.1 μM (NH4)6Mo7O24, and 50 μM KCl, in presence of 1% (wt/vol) sucrose and 0.8% (wt/vol) agar. -Zn or -Fe medium was made by not adding the only source of Zn (ZnSO_4_) or FeEDTA to the medium, respectively. For the assays involving azelaic acid treatments, AzA (246379 ALDRICH, Sigma) was added in different concentration ranging from 25 to 200 µM. For the assay with rice (*Oryza sativa* L.), Niponbare was used and seeds were soaked in deionized water over night in dark then transferred in a controlled-environment chamber (light/dark cycle of 14/10 h, 200 μmol photons m^-2^s^-1^, temperature of 28/25 °C and RH of 80%) to ¼ Yoshida media for 5 d ^50,51^: 0.36 mM NH_4_NO_3_; 0.41 mM MgSO_4_; 0.19 mM CaCl_2_; 0.13 mM K_2_SO_4_; 0.08 mM NaH_2_PO_4_; 4.72 µM H_3_BO_3_; 2.37 µM MnCl_2_; 8.90 µM Fe-NaEDTA; 0.62 µM ZnSO_4_; 0.04 µM CuSO_4_; 0.02 µM (NH_4_)_6_Mo_7_O_24_, adjust to pH 5.5. ZnSO4 was removed in -Zn medium. For rice treatments, azelaic acid (246379 ALDRICH, Sigma) was added in different concentration ranging from 75 to 300 µM. For the control condition, rice plants were kept in nutrient solution with the above-mentioned composition. Rice seedlings were grown in a growth chamber under the following environmental conditions: light/dark cycle of 14/10 h, temperature of 28/25 °C, and RH of 80 %.

GWAS root trait quantification was conducted using the BRAT software^38^. Root length in assays involving mutant and transgenic lines, as well as rice plants was measured using ImageJ software, version 2.0.0 (http://rsb.info.nih.gov/ij/). Statistical differences between genotypes were calculated using t-test analyses and ANOVA with subsequent post hoc tests using Graphpad Prism (GraphPad Software Inc., San Diego, CA, USA) or Microsoft Excel (Microsoft, USA).

### Gene expression analysis by quantitative RT-PCR

Total RNA was extracted from roots of Arabidopsis wild type plants (different accessions) grown in presence or absence of Zn, and with or without azelaic acid (246379 ALDRICH, Sigma), using Plant RNeasy extraction kit (Qiagen) and RQ1 RNAse-free DNAse (Promega). Two μ g of total RNA were used to synthesize cDNA using poly-A oligos. Real-time-quantitative reverse-transcription PCR (RT-qPCR) was performed with a Light Cycler 480 Real-Time PCR System (Roche; Roche Diagnostics, Basel, Switzerland) using LightCycler 480 SYBR Green I Master mix (Roche, IN, USA). Primer list is provided in table S6. Gene transcript accumulation quantification were performed in a final volume of 20 μL containing optimal primer concentration 0,3 µmol, 10 μL of the SYBR Green I master mix, and 5 μL of a 1:25 cDNA dilution. Real time-PCR conditions were as 95°C for 5 min, and followed by 40 cycles of 95°C for 10 s, 60°C for 10 s, 72 °C for 25 s, and finally one cycle 72 °C for 5 min. As a negative control, template cDNA was replaced by water. All PCR reactions were performed in triplicates. For each sample, cycle threshold a (Ct) value was calculated from the amplification curves. For each gene, the relative amount of calculated mRNA was normalized to the level of the control gene *ubiquitin10* mRNA (*UBQ10*: At4g05320). For every sample, the relative gene expression of each genes was expressed following normalization against the CT values obtained for the gene used for standardization, for instance Δ CT,*AZI1* = CT,*AZI1* − (CT,*UBQ10*). Quantification of the relative transcript levels was performed as described previously^52,53^. Briefly, -Zn treatment was compared to +Zn, the relative expression of a each gene was expressed as a ΔΔ Ct value calculated as follows: ΔΔ Ct = Δ CT,*AZI1*(-Zn) – Δ CT, *AZI1*(+Zn). The fold change in relative gene expression was determined as 2− ΔΔ CT.

### Bacterial strains and growth conditions

Wild-type strain of *Azospirillum brasilense* is used in this study^54^. These bacteria strains were cultivated and inoculated in plant culture medium as described previously^55^. Statistical differences between genotypes were calculated using t-test analyses and ANOVA with subsequent post hoc tests using Graphpad Prism (GraphPad Software Inc., San Diego, CA, USA).

### Plasmid construction and plant transformation

The *AZI1* locus from Col-0 and Sq-1 accessions were cloned with primers spanning the region ranging from 1614 bp upstream of the *AZI1* transcription start site to the stop codon of *AZI1* into the binary vector pCAMBIA1301 by restriction enzymes of *BamHI* and *PstI* using primers listed in Supplementary table S3 and S4. The constructs were transformed into *Agrobacterium tumefaciens* strain GV3101 and then used for Arabidopsis transformation by the floral dip method^56^. Transgenic plants were selected by antibiotic resistance, and only homozygous descendants of heterozygous T2 plants segregating 1:3 for antibiotic resistance: sensitivity were used for analysis.

### GWA mapping

For GWAS, mean total root length values of 230 natural accessions were used (Table S1, S2). The GWA analysis was performed in the GWAPP web interface using the mixed model algorithm (AMM) that accounts for population structure^26^ and using the SNP data from the RegMap panel^57,58,22^. Only SNPs with minor allele counts greater or equal to 10 were taken into account. The significance of SNP associations was determined at 10% FDR threshold computed by the Benjamini-Hochberg-Yekutieli method to correct for multiple testing ^27^.

---

**Supplementary Table 1.** Primary root growth of 230 natural accessions of *Arabidopsis thaliana* grown on zinc sufficient (+Zn) medium over 7 days after germination.

**Supplementary Table 2.** Primary root growth of 230 natural accessions of *Arabidopsis thaliana* grown on Zn limiting conditions (-Zn) over 7 days after germination.

**Supplementary Table 3.** Broad sense heritabilities for growth under –Zn conditions.

**Supplementary Table 4.** Mean of primary root length of accessions measured at day2 and *AZI1* locus marker SNP-allele (Chr4 7400493) of all accessions used in this work.

**Supplementary Table 5.** List of primers used in this study.

**Figure S1. Genetic diversity and genotype by -Zn dependent root growth responses.** A) Genetic diversity of accessions used in this study. Plotted are the two major principal components (PCs) from^22^that infer continuous axes of genetic variation. B, C) Mean root lengths (pixels) of accessions grown in +Zn (x-axis) or -Zn (y-axis) growth conditions on the days for which significant GWAS signals were found. Day 2 (B); Day 7(C).

**Figure S2. mRNA abundance of Zn-responsive genes ZIP3, ZIP5, ZIP12 and PHO1;H3 in roots of Col-0 plants exposed to different Zn availabilities.** Transcript levels of *ZIP3* (At2g32270), *ZIP5* (At1g05300), *ZIP12* (At5g62160) and *PHO1;H3* (At1g14040) in roots of Arabidopsis (Col-0) seedlings grown on vertical agar plate in presence or absence of Zn (day 5) as determined by Real-time qPCR. Transcript levels of these genes are expressed relative to the average transcript abundance of *UBQ10* (At4g05320) that was used as an internal control, and relative to +Zn values that were set to 1. Every data point was obtained from the analysis of roots collected from a pool of six plants. Data presented are means of three biological replicates ± SE. Asterisks indicate statistically significant differences compared to the +Zn treatment for each gene (P <0.05).

**Figure S3. Frequency distribution of mean primary root length of Arabidopsis accessions grown on -Zn.** Histograms of the daily frequency distribution of mean primary root length of 230 Arabidopsis accessions under -Zn conditions over a time course of seven days.

**Figure S4. GWASs of mean root length grown on +Zn.** Manhattan Plots showing the genome-wide associations of mean root length in a set of 230 *A. thaliana* accessions grown in presence of zinc for seven days. The chromosomes are represented in different colours. The horizontal blue dash-dot line corresponds to a nominal 0.1 significance threshold after Benjamini–Hochberg–Yekutieli correction.

**Figure S5. GWASs of mean root length grown on –Zn.** Manhattan Plots showing the genome-wide associations of mean root length in a set of 230 *A. thaliana* accessions grown under Zn limiting conditions for seven days. The chromosomes are represented in different colours. The horizontal blue dash-dot line corresponds to a nominal 0.1 significance threshold after Benjamini–Hochberg–Yekutieli correction.

**Figure S6. Primary root length of azi1 mutant, 35S AZI1 (OE AZI), Col-0 plants grown on +Zn and -Zn.** Average primary root length of wild-type plants (Col-0 genotype), *azi1* mutant and overexpressor line 35S::*AZI1* (OE *AZI1*) plants grown under +Zn or –Zn over a time course of seven days. Experiments were independently repeated three times, and data are represented as mean ± s.d. n = 10. Letters a, b and c indicate significantly different values at p <0.05 determined by one-way ANOVA and Tukey HSD.

**Figure S7. AZI1 is not involved in control of root growth under Zn or Fe limited conditions.** (A) Representative root growth phenotypes of seedling (day 5) grown under +Zn or –Zn conditions. Shown are wild-type plants (Col-0 genotype), azi1 mutant and overxpressor line (OE *AZI1*) 35S::*AZI1* plants. (B) Average primary root length of wild-type plants (Col-0 genotype), azi1 mutant and overexpressor line (OE *AZI1*) 35S::*AZI1* plants, day 5, grown under +Zn or –Zn respectively. (C) Average primary root length of wild-type (Col-0), azi1 mutant and overexpressor line 35S::*AZI1* (OE *AZI1*) plants, day 5, grown under +Fe or –Fe, respectively. Experiments were independently repeated three times, and data are represented as mean ± s.d. n = 10. Letters indicate significantly different values at p <0.05 determined by one-way ANOVA and Tukey HSD.

**Figure S8. Polymorphism patterns around the AZI1 locus in extreme accessions.** Gene models and SNP polymorphisms among representative extreme accessions (4 accessions with short root phenotype and 4 accessions with long root phenotype for the genomic region surrounding the *AZI1* gene. (A) Amino acid changes around *AZI1* (At4g12470) locus. (B) SNPs around *AZI1* locus; Synonymous amino acid: green line, non-synonymous amino acid: red line. Only genomes that were available in the SALK 1001 genomes browser (http://signal.salk.edu/atg1001/3.0/gebrowser.php) as of August 2016 were considered. (C) Representative images of contrasting PRG phenotype (day 5) of eight Arabidopsis thaliana accessions grown in +Zn or –Zn conditions, scale bars: 1 cm. (D) Relative transcripts accumulation of AZI1 gene in these eight accessions grown in –Zn conditions compared to +Zn conditions. Arabidopsis *Ubiquitin* gene was used as an internal reference. The data are given as means ± s.d. n = 10.

**Figure S9. Natural allelic variation of the AZI1 locus is not relevant for root length variation in Zn sufficient condition.** Primary root length of *azi1* lines complemented with pAZI1*:AZI* from either Col-0 (n=44), Sq-1 (n=44) or the empty vector (n=27) on +Zn conditions for five days. For each genotype, three repeats each containing five independent T3 lines. Box plots show analysis of the primary root length of *azi1* lines complemented with pAZI1:*AZI* from either Col-0, Sq-1 or the empty vector on +Zn conditions. Center lines show the medians; box limits indicate the 25th and 75th percentiles as determined by R software; whiskers extend 1.5 times the interquartile range from the 25th and 75th percentiles.

**Figure S10. Alignment of the promoters of Arabidopsis AZI1 genes from Col-0 and Sq-1 accessions.** Alignment was done using the MultAlign program (http://bioinfo.genopole-toulouse.prd.fr/multalin/multalin.html).

**Figure S11. Expression levels of defence-related genes.** Transcripts accumulation of At2g31890, At5g54250, At5g24150, At1g09070, At5g60890, At1g76850, At4g04210, At3g54840, At5g66760, At2g37190, At1g14980, At1g10585, *WRKY46* (At2g46400), At3g12580, *AZI1* (At4g12470), *ICS1* (At1g74710), At5g47230, *WRKY28* (At4g18170), At2g18210, At1g27730, At5g20030 and *PR1* (AT2G14610) in *Arabidopsis thaliana* Col-0 genotype grown in +Zn or –Zn conditions with or without 100µM AzA. The Arabidopsis *Ubiquitin* gene was used as an internal reference. Experiments were independently repeated three times. Data was normalized to +Zn/-AzA condition, log2 transformed and visualized using "clustered correlation" CIM (http://discover.nci.nih.gov)^59^. Stars indicate genes involved in the biosynthesis and response to salicylic acid accumulation.

**Figure S12. Conservation of the Zn/AzA interaction in rice.** Rice (*Oryza sativa*, Nipponbare) seedling were grown for five days starting the day of imbibition on Yoshida medium supplemented with 300 µM of AzA in presence or absence of Zn. Scans were taken from 5-days old seedlings. Red arrows indicate roots.

